# Water fluxes contribute to growth patterning in shoot meristems

**DOI:** 10.1101/2023.08.27.554993

**Authors:** Juan Alonso-Serra, Ibrahim Cheddadi, Annamaria Kiss, Guillaume Cerutti, Claire Lionnet, Christophe Godin, Olivier Hamant

## Abstract

In multicellular organisms, localized tissue outgrowth creates a new water sink thereby modifying hydraulic patterns at the organ level. These fluxes are often considered passive by-products of development and their patterning and potential contribution to morphogenesis remains largely unexplored. Here, we generated a complete map of cell volumetric growth and deformation across the shoot apex in *Arabidopsis thaliana*. Within the organ-meristem boundary, we found that a subpopulation of cells next to fast-growing cells experiences volumetric shrinkage. To understand this process, we used a vertex-based model integrating mechanics and hydraulics, informed by the measured growth rates. Organ outgrowth simulations revealed the emerging water fluxes and predicted water deficit with volume loss for a few cells at the boundary. Consistently, *in planta,* a water-soluble dye is preferentially allocated to fast-growing tissues and fails to enter the boundary domain. Analysis of intact meristems further validated our model by revealing cell shrinkage next to fast-growing cells in different contexts of tissue surface curvature and cell deformation. A molecular signature of water deficit at the boundary further confirmed our conclusion. Taken together, we propose a model where the differential sink strength of emerging organs prescribes the hydraulic patterns that define the boundary domain at the shoot apex.

## Introduction

During development, multicellular tissues are patterned by biochemical and mechanical cues (1)(2)(3)(4)(5)(6)(7). This is notably the case at the shoot apical meristem (SAM) which contains the plant stem cell niche and where hormone and peptide patterns define the different zones and their function: self-maintenance (central zone (CZ)), organ production and identity (peripheral zone (PZ)) and organ separation and axillary meristem initiation (boundary zone (BZ)). In particular, differential growth together with tissue shape prescribe a stereotypical pattern of mechanical stress, impacting cytoskeleton behavior (8), hormone carrier polarity (9)(10), chromatin status (11) and gene expression (12) in a feedback loop. Other physical cues, such as hypoxia (13), have been integrated in this picture more recently. However, until now, the role of water has been considered non-limiting for morphogenesis, with water playing essentially no role in the patterning *per se*.

In the SAM, organs develop sequentially following a robust spatiotemporal organization, but vascularization is temporally decoupled from early organ growth so water must necessarily travel from the vasculature through non-vascularized parenchymatic and meristematic cells (14). In most aboveground plant tissues water movement is driven by stomata transpiration, but shoot meristems lack stomata and water is expected to adapt to growing organs and fill them. Interestingly, growth *per se* has the capacity of lowering the water potential, thereby attracting water. It has been proposed as an additional driver of water movement in plants (15)(16). This is what we investigate here, taking advantage of the plant meristem as a model system.

First, we obtained a comprehensive map of volumetric growth in the inflorescence SAM including developing organs up to stage 2 (before the flower organs appear). In addition to the expected differential growth rate between the organs and the meristem, we found evidence of cell shrinkage specifically in the boundary domain. Using a modeling approach informed by our morphometric analysis we predicted that higher growth rates in the organ may affect the hydraulic pattern and cause water deficit in the adjacent boundary tissue. The allocation pattern of a water-soluble dye was in agreement with our model and revealed a degree of symplastic flow isolation in the boundary domain. Finally, by inducing *de novo* organ growth in intact meristems we found further evidence of cell shrinkage adjacent to fast-growing cells.

Altogether, our study demonstrates that water is a factor whose non-uniform flux distribution contributes to the pattern of tissue development, in synergy with biochemical and mechanical cues.

## Results

### Global volumetric analysis of the SAM reveals shrinkage of deep boundary cells

Previous morphometric analyses of shoot meristems have revealed the cell volumetric growth dynamics of the CZ (17) or the initial stages of flower development (18). Yet, these quantifications have not been related to water fluxes and have not been extended to later developmental stages with highly folded tissues. Growth and deformation rates of the area of the outer periclinal wall have been described for complete meristems of pimpernel, tomato, and Arabidopsis (19)(20), but without volumetric insights. In order to understand the growth-driven hydraulic patterns in the whole SAM while considering tissue deformation we measured cell volumetric changes along with tissue topology, and changes in cell sphericity, during organ emergence. To do so, we performed time-lapse acquisitions using a plasma membrane marker and 3D watershed segmentation using timagetk tools (Fig. 1A, S1; see Methods).

**Fig. 1.**
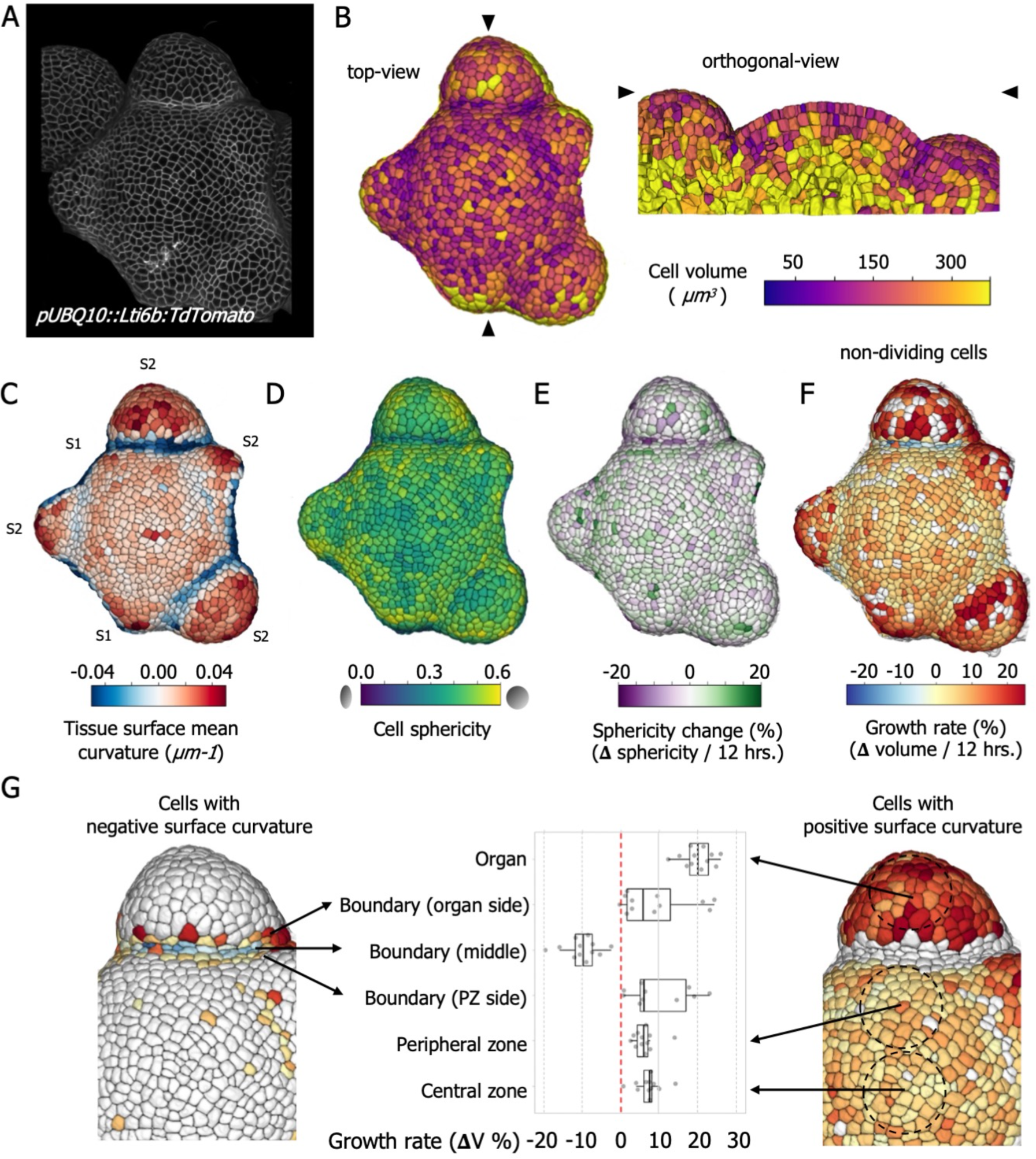
**Morphometric analysis of the SAM reveals volumetric shrinkage at the deep boundary.** (A) Processed signal of *pUBQ10::LTi6B-TdTomato* marker used for 3D segmentation (L1 only shown). (B) Heatmaps showing cell volumes from the top view (L1 only) and from an orthogonal view obtained on the site of the black arrows. (C) Heatmaps showing the surface mean curvature and the organ stages 1 (S1) and 2 (S2). (D) Heatmap showing cell sphericity. (E) Heatmap showing the sphericity changes over 12 hours. (F) Heatmap showing volumetric growth rates over 12 hours. Dividing cells appear in white so growth rates are only shown for non-dividing cells. (G) Growth rate maps of a representative organ-meristem region including dividing cells (same sample as in panel F). The volume of daughter cells was added to obtain tissue-level growth rate. Note in panel F that no cell division was detected in the boundary. The subset of cells with negative curvature (left) and positive curvature (right). The boxplots in the middle reveals the position-dependent differential growth rates of cells for one meristem. The same heatmap profile was detected in all studied meristems (n=6).

The cell volume distribution was heterogenous with larger cells in developing organs and inner layers below the third layer of cells (L3) (Fig. 1B). The tissue surface mean curvature describes the topography of the meristem and serves as a morphological marker to identify the organ stage. Organs in stage 1 do not exhibit a morphological crease at the boundary (21) whereas in stage 2-organs the saddle-shaped surface becomes significant (defined when the amplitude of the negative curvature is higher than that of the maximum curvature) (Fig. 1C). While cell sphericity and sphericity changes are variable in time and space, boundary cells are progressively deformed and become markedly least spherical in stage 2-organs (Fig. 1D, E).

Then we quantified the volumetric changes, selecting cells that did not undergo division over 12 hours. As expected, the younger organs and early boundary cells exhibited positive growth rates in this time window (Fig. 1F). In early boundaries, the mean curvature was positive and boundary cells, grew by 6.9 ± 3.8% while organ cells (stage 1) grew by 18.2 ± 6.8%. When the primordium reaches stage 2, the mean curvature became negative in the boundary. In this organ stage, more cell divisions were detected and cells had a higher volumetric growth rate (Fig. 1F). However, at later stages, i.e. three plastochrons after the transition to stage 2 (from primordium 5 (P5) onwards), we observed a new pattern at the boundary: cells localized either on the organ or on the PZ sides were expanding, while cells localized in the deep boundary domain were shrinking (Fig. 1G). This stage also coincided with a loss of sphericity in boundary cells.

The shrinkage was of significant amplitude: shrinking cells lost on average 10.1 ± 4.6 % of their volume over 12 hours, while the adjacent subpopulations (with negative mean curvature too) exhibited a highly heterogenous expansion rate of 9.7 ± 9% on the organ side and 9.8 ± 8% on the PZ side. In the same time window, cells with positive mean curvature at the top of the organ, grew by 19.9 ± 4 % and cells in the PZ 6.1 ± 3 % and CZ by 7.2 ± 3.3 %. This pattern was found in 6 meristems from 6 independent plants (Fig. S2A). We also estimated the measurement error due to a different resolution in XY vs. Z. A multiangle analysis revealed an average of 0.3% error in volume estimation due to Z differential resolution (Fig. S7B,C).

Including dividing cells to the growth map did not change the trend (Fig. 1F,G), consistent with the fact that cells in late boundaries rarely divide (22), and that most dividing cells contribute to organ and PZ expanding tissues. This analysis reveals that late boundary cells, despite being confined to a narrow space, can display opposing growth behaviors. A few of them lose volume without division, potentially revealing the consequences of growth-driven hydraulic patterns in the SAM.

### Water efflux from the deep boundary domain is predicted by a mechano-hydraulic model

The above analysis suggests that boundary cells could experience a local water deficit as a consequence of the volume loss. This echoes a previous computational study (23), which suggested that, in theory, local tissue growth could induce water loss in cells surrounding the growing region. To explore this hypothesis in our context, we adapted our 2D-multicellular model (23) to capture the key organization of cells in a SAM section, including the central zone, peripheral zone, boundaries, and primordium (Fig. 2, Fig. S3A). The model is a multicellular extension of the Lockhart-Ortega model (24)(Fig.2A-C), it assumes that every cell has a water potential composed of i) a constant osmotic potential and ii) a varying hydrostatic pressure. Water fluxes follow decreasing water potential; influx or efflux of water induces cell volume change (Fig. 2B), and mechanical constraints in cell walls accommodate for it (Fig. 2C). This, in turn, impacts the cell hydrostatic pressure (Fig. 2A), which creates a complex feedback loop between cell water potential, fluxes, and mechanical stresses in cell.

**Fig. 2.**
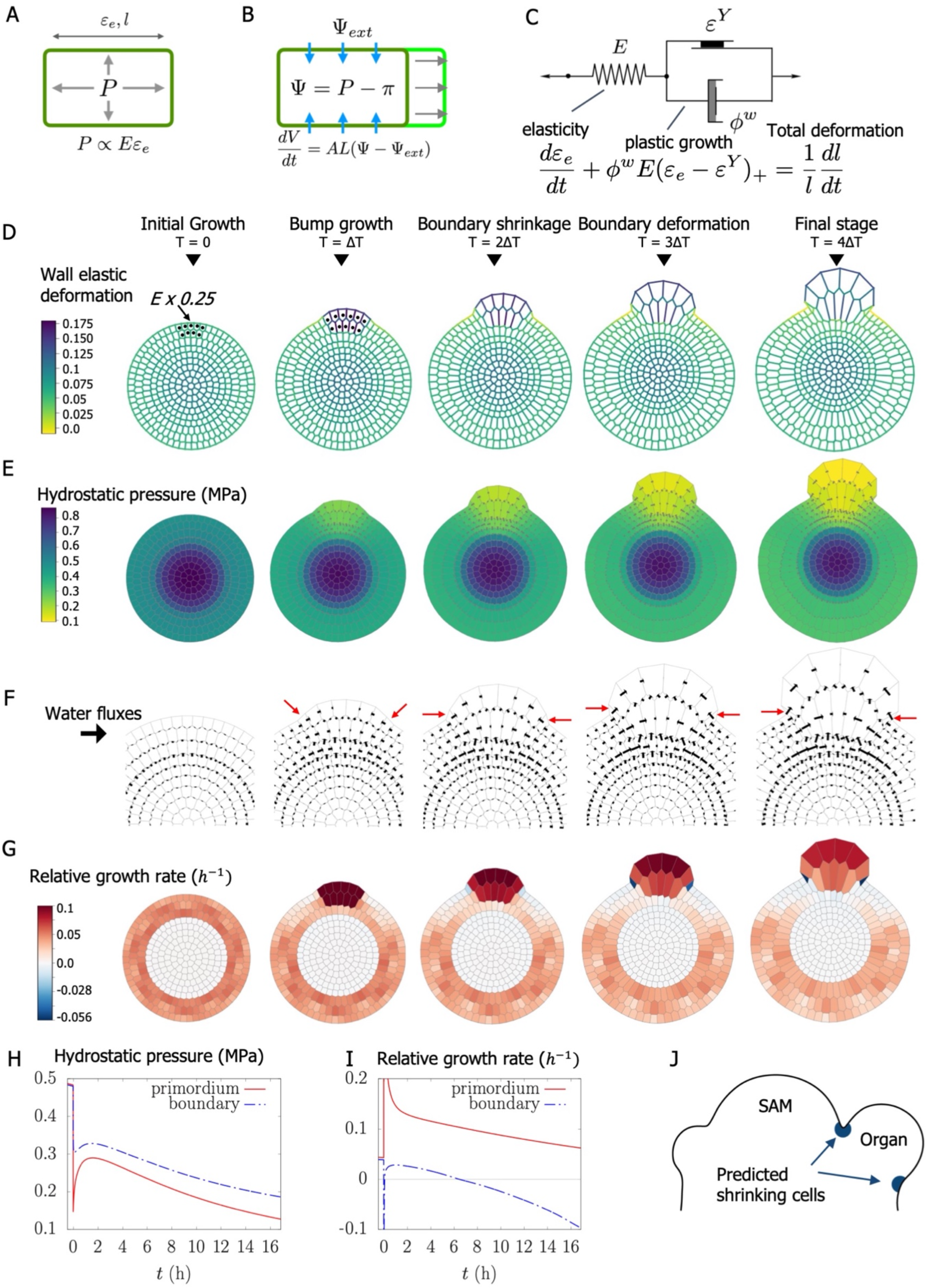
**Vertex-based model of growth-derived water fluxes.** (A-C) Lockhart-Ortega elongating single-cell model: (A) Cell pressure and wall tension equilibrate: pressure ***P*** is proportional to elastic modulus ***E*** and elastic deformation ***ε_e_***. (B) Volume variation and water fluxes over surface ***A*** of permeability ***L*** depend on the difference between cell (**Ψ**) and external (**Ψ**_***ext***_) water potentials. (C) Wall mechanical response decomposes into elastic (1^st^ term in left-hand side) and plastic (2^nd^ term) responses; the plastic response corresponds to irreversible growth characterized by extensibility ***ϕ^w^*** and is triggered when the elastic deformation is above the threshold ***ε^Y^***. The driving force is a large enough value of osmotic pressure ***π*** that induces water flux into the cell, which triggers simultaneously volume increase, wall deformation, pressure, and growth. (D-G) Multicellular model at regular intervals **Δ*T* = 4. 3 *h***, from the initial condition to the final stage: cells in the center deform elastically but cannot grow, they accumulate water from an external source and pressure; conversely, wall growth relaxes tension in cells in L1-L3; thanks to their lower pressure, they attract water from the center. To mimic organ outgrowth, *E* is reduced (*E* x 0.25) in a subgroup of L1L2 cells (black dots) at time ***T*** = **0**. (D-G) Maps of elastic deformation, hydrostatic pressure, fluxes, and relative growth rate; the boundary cells are indicated by red arrows in (F). (H-I) Time evolution of pressure (H) and relative growth rate (I) in boundary cells (red arrows in (F)) and in the primordium (mean value). Pressure and growth rate global decrease is an effect of the global growth of the tissue (see Supp. Info). (J) Cartoon model of an inflorescence SAM showing the sectors where cell shrinkage is predicted.

The model parameters are adjusted such that L1, L2, and L3 cells are growing thanks to the water provided by cells underneath (Fig. S3A, Table S1). The initiation of a primordium outgrowth is triggered by a decrease of the elastic modulus in the cell walls of the future organ (Fig. 2D). Importantly, in the current model we adjusted the organ elastic modulus such that organ cells would grow 4 times faster than the rest of the tissue as indicated by our morphometric measurements (see Fig. 1G). Simultaneously, wall deformation increases, and wall stresses and cell hydrostatic pressure decrease in the initiated organ (Fig. 2E, H and Fig. S3A, B). This local drop in pressure in the growing organ creates a water sink that draws water from neighboring cells (Fig. 2F). Water movement appears as an emerging property and fluxes are rapidly polarized to feed the growing organ. Hence, the boundary cells (indicated by red arrows in Fig. 2F) are caught between peripheral zone cells and organ cells, with limited access to water on one side and high demand on the other (red arrows in Fig. 2F). In the first stage, they are hardly able to maintain positive growth. In the second stage, the water demand increases as the organ grows, up to the point where boundary cells begin to lose volume at a significant rate (Fig. 2 G, I and Fig. S3 D, E).

To assess the robustness of this shrinkage effect in different growth conditions, we varied both the wall stiffness at the time of organ outgrowth (which affects growth rates) and the cell-to-cell water conductivity across L1L2L3 layers. Simulations resulted in different topography and growth rate dynamics of the primordium (Fig. S4). Yet, the boundary-specific cell shrinkage was observed in all tested scenarios, suggesting that lateral water depletion is a robust emerging property of local primordium outgrowth.

### A water-soluble dye does not reach the deep boundary domain

To check whether the deep boundary would be the site of such water depletion, we explored gene expression across different SAM domains using an available translatome from vegetative meristems (25), reasoning that a global molecular signature may provide indirect evidence of water loss. By performing a GO enrichment analysis on each subset of domain-specific upregulated genes, we found that genes associated to “abiotic and biotic stress” are enriched in the WUS, LAS and PTL domains (Data S1). The LAS domain, which best matches the early to late boundary domain, was differentially enriched for genes associated to “response to osmotic stress” and “response to salt stress” (Fig. S5A). This is consistent with local water depletion in that domain.

To get closer to causality, we would need to monitor water movements in our system. Water being in essence invisible under an optical microscope, its behavior can only be inferred with dyes, as a proxy for water movement. HPTS (8-hydroxypyrene 1,3,6 trisulfonic acid) is a membrane-impermeable, water-soluble dye, that has previously been used to reveal leaf-to-apex vascular transport and symplastic domains in vegetative and inflorescence shoot apices (26). Here we used a plasma membrane marker to resolve the anatomy in internal layers in the SAM and provided HPTS dye such that it would be acropetally transported from the stem. Results presented below were obtained in 10 meristems from 10 independent plants.

After 1, 3, 6, 12 or 24 hours of HPTS incubation, we detected dye signal in the SAM (Fig. 3A,B and S5C). However, pattern and signal intensity were more consistent across samples after 6 and 12 hours of incubation. After 6 hours of incubation, and in line with a previous report (26), the HPTS signal was enriched in L1, L2, and L3 with a bias towards developing organs (Fig. 3C). A longer incubation time of 12 hours further highlights an organ-enriched pattern with HPTS dye being present in the cytoplasm and concentrated in vacuoles even in young developing primordia (P1 to P5) (Fig. 3B and S5B). Furthermore, while longer HPTS incubation times of up to 24 hours increased the overall signal intensity, the described pattern remained the same (Fig. S5C). This pattern suggests that organs are indeed preferential sinks for symplastic fluxes.

**Fig 3.**
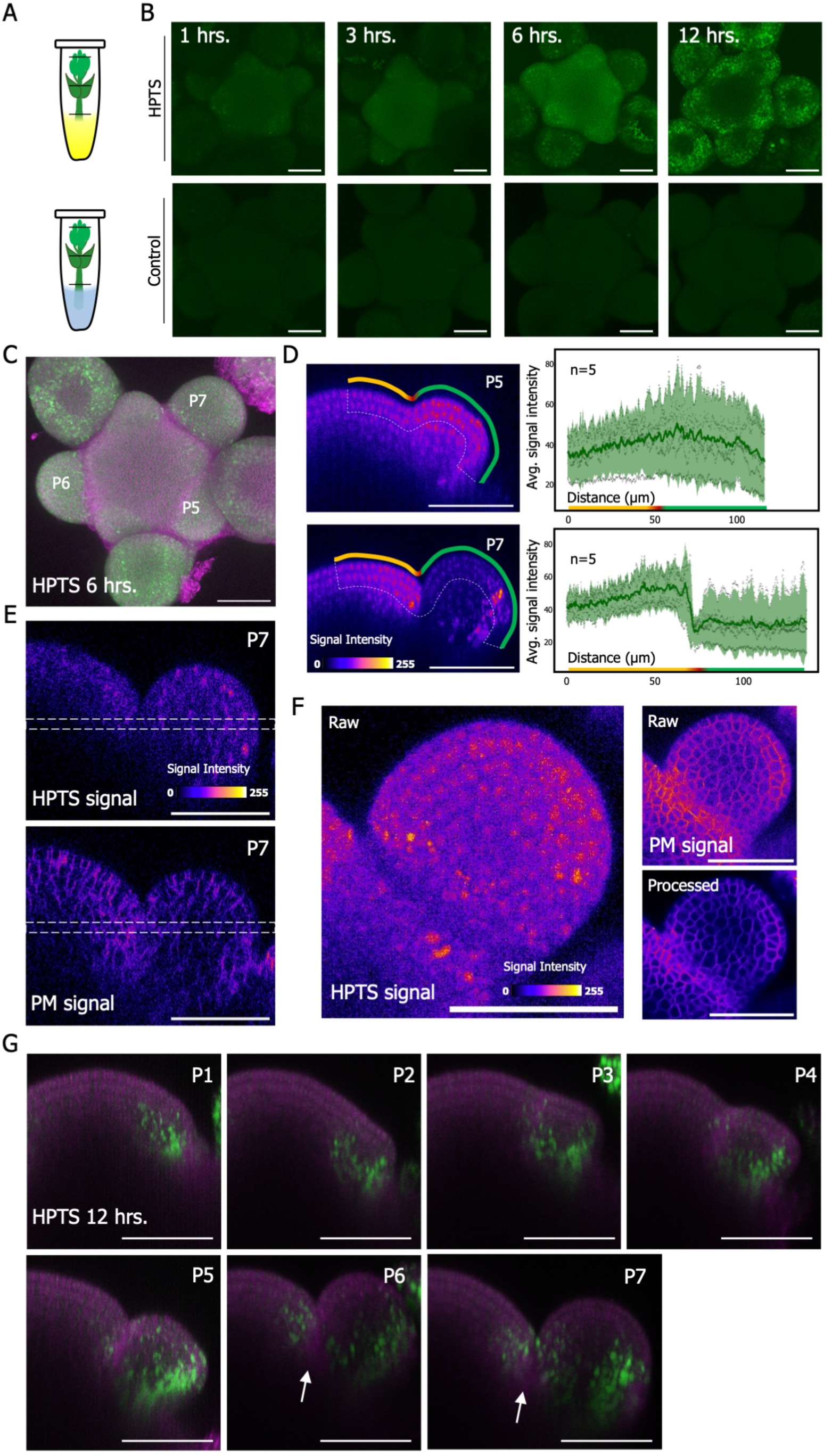
**HPTS allocation patterns highlight the differential sink strength of organs.** (A) Drawing showing the experimental setup. Shoot apices were treated by dipping stems in liquid ACM media with (top) or without HPTS (bottom). (B) HPTS (green) pattern after different incubation times in dissected meristems. (C) Maximum intensity projection of HPTS pattern with the plasma membrane marker after 6 hours of incubation. (D) HPTS signal intensity profile plot showing the average (green line) and 95% confidence interval (light green) for L1L2 extracted from orthogonal sections of P5 and P7 after 6 hours of dye incubation (n=5). Orthogonal sections illustrate the sampled layers (dashed white line in L1 and L2) across meristem domains: top yellow line (CZ and PZ), red line (BZ) and green line (organ), see methods for sampling details. (E) Orthogonal sections showing HPTS and plasma membrane (*pUBQ10::LTi6B-TdTomato*) signal intensities for P7 organs after 6 hours of incubation (stacks from panel C). The white dotted lines define the limits of the stack of slices that were projected to obtain panel F. (F) Maximal projection of a 5*μ*m-thick transversal section in P7 organs showing raw data (HPTS) or raw and anisotropic filtered images for better resolution of plasma membrane signal. Signal intensity is color-coded. (G) Orthogonal sections showing organ stage-dependent HPTS pattern (as defined in Fig. S5B) after 12 hours of incubation. Deep boundary domains with reduced HPTS levels are marked with white arrows.

In order to characterize the earliest HPTS allocation pattern we further quantified the HPTS signal after just 6 hours of incubation. We detected a gradient of increasing HPTS intensity from the meristem to P5 stage organs (Fig. 3D). Strikingly, the boundary of advanced organs (P6 to P7) displayed a marked drop in HPTS signal compared to the meristem or the organ, as viewed from the side or above (Fig. 3D-F and S5D). Note that the signal of ubiquitously expressed plasma membrane marker did not exhibit such a pattern, indicating that the HPTS depletion zone is not an optical artefact caused by tissue depth or curvature (Fig. 3F). Thus, HPTS is excluded from the boundary domain in that time period.

To check whether HPTS could ultimately reach the boundary domain, we monitored its distribution using increased resolution and correlative imaging with a membrane marker. After 12 hours, HPTS accumulated in L2 and L3 of early-stage flower primordia (before bulging out) and flower organs (emerging outer sepal) (Fig. 3G). We also observed an asymmetric signal distribution in stage 2 organs (e.g. P5 to P7), with higher signal on abaxial side (Fig. S5D), consistent with increased growth rate in that domain (18). Yet, HPTS was still absent from the boundary domain (Fig. 3G, white arrows).

To conclude, while water is ubiquitous to all cells in the SAM, the differential sink strength of organs creates a HPTS allocation pattern that is in agreement with our growth-induced water flux model.

### Differential growth rates generate cell shrinking domains in intact plants

In the previous experiments, shoot apices were cut from the stem and most organs were removed to facilitate confocal imaging. To control for potential dissection-induced hydraulic artefacts on cell growth, we repeated the analysis using intact plants. To this end, we pre-treated *in vitro*-growing plants with N-1-naphthylphthalamic acid (NPA), which results in the development of pin-like shoot apices, fully accessible to optical microscopy. 48 hours after transfer of the whole plant to an NPA-free medium, the meristem recovers and starts generating organs, allowing confocal access without organ dissection or stem cutting (27)(see Methods).

In this system the growth rates were 4 to 6 times faster than in meristems from dissected shoot apices, thus requiring more frequent image acquisitions to capture the boundary formation.

Tissue surface curvature and cell sphericity changes similar to those observed before were detected in just 2 to 4 hours (Fig. 4 A,B). The volumetric analysis revealed that organ cells grow by more than 20% (Fig. 4 C,D). In agreement with our previous observations, we detected shrinking cells losing 7-10% of their volume in the midline and flanks of the newly formed boundary (Fig. 4C-D; n=6). This confirms that a subpopulation of late boundary cells shrinks, consistent with predicted water efflux simulations, independent of dissection.

**Fig 4.**
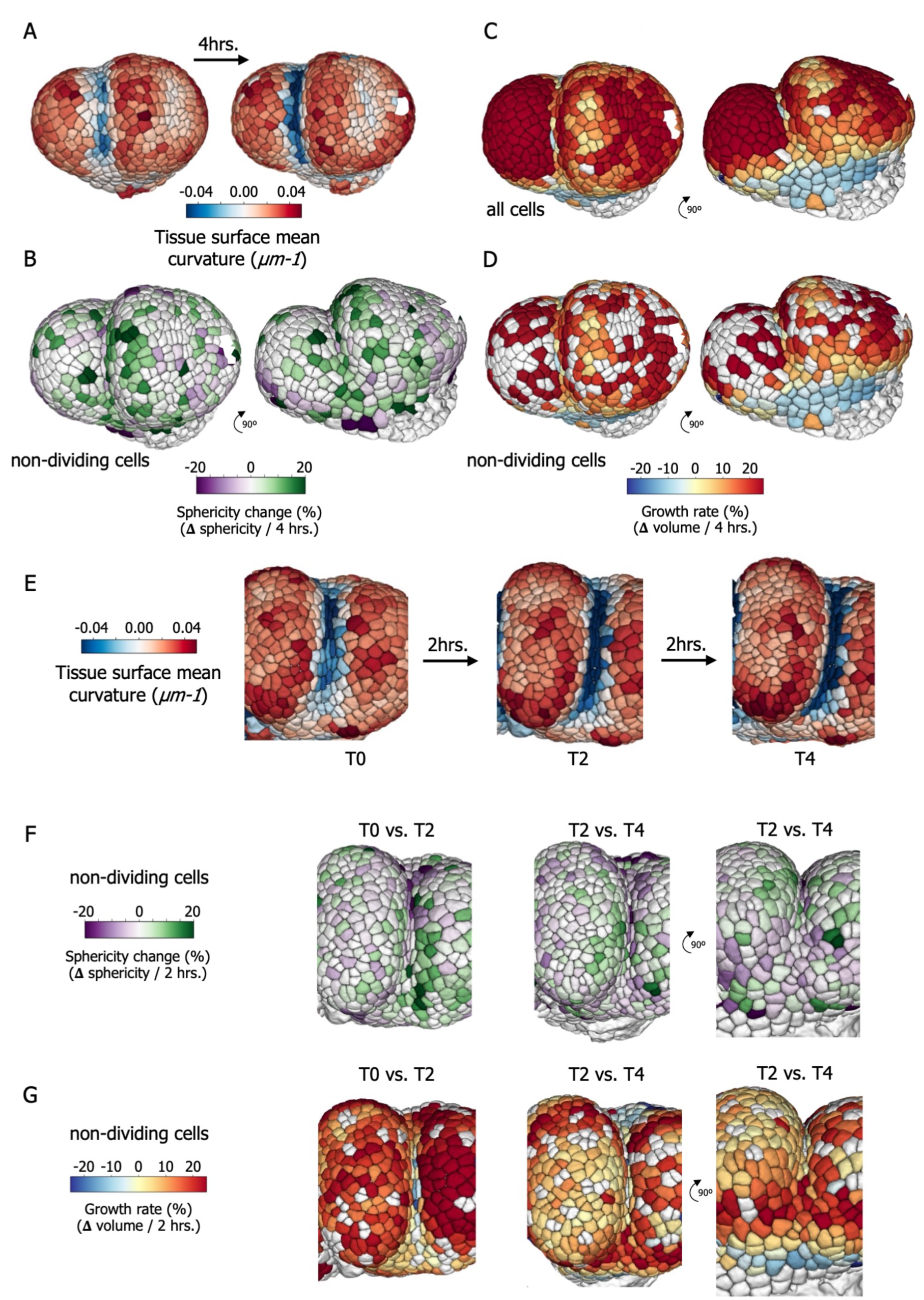
**Negative growth is observed in deep boundaries from intact (non-dissected) pin-like shoot apices** (A) Early organ outgrowth heatmap showing the surface curvature before and after 4 hrs. (B) Heatmap showing the cell sphericity changes during the same 4 hrs. of growth. (C) Heatmaps represent the volumetric growth rates including cell divisions (C) or marking dividing cells in white (D). The same samples were rotated 90° to visualize shrinking cells detected at the boundary flanking sides. (E) Three time points show the developmental progression of an advanced boundary with heatmaps representing the surface mean curvature (E), and the corresponding cell sphericity changes (F) and volumetric growth rates (G) for non-dividing cells. Note the presence of pale blue cells, i.e. shrinking cells, in deep boundary (left image) and below the meristem (right image).

Strikingly, we also observed the presence of many epidermal cells shrinking below the meristem (Fig. 4D,G). Unlike cells in the midline boundary, the surface mean curvature for cells below the meristem was less pronounced or even positive, and not all of them showed loss of sphericity (Fig. 4E,F). This suggests that shrinkage could be more widespread than expected, especially in cells neighboring tissues with very fast growth. Altogether, we systematically detected decreases in cell volume in boundary regions, both in dissected and non-dissected meristems, which supports the water loss hypothesis in these cells.

## Discussion

In essence, morphogenesis entails a change in shape, and thus a change in structure, i.e. mechanics. In turn, mechanical stress together with molecular patterning factors control morphogenesis. Here we propose that growth-induced deformation not only impacts the local stress pattern, but it also affects the hydraulic pattern, which in turn helps to define boundaries.

Note that one specificity of the shoot apex is that the water-draining role of the growing organ acts in synergy with tissue folding: the deep boundary cells are locked in a domain where tissue deformation geometrically restricts cell expansion. This might explain why differences in volumes are so significant in shrinking cells, and why these cells exhibit unusual behavior (reduced division rate (22), specific gene expression pattern (28)) and fate (29).

Even in narrow domains of the SAM, hydraulic properties such as turgor pressure are heterogenous and depend e.g. on tissue topology (31). Our complementary finding of shrinking domains next to fast-growing sectors supports the long-proposed hypothesis of growth-induced water potentials and water transport across the tissue (15). Thus, late boundaries shrinkage is partly a hydraulic consequence of organ outgrowth, which contributes to the final tissue topography. This makes water a patterning factor at the shoot apex acting in synergy with mechanical and biochemical cues.

Taken together, our results open a new avenue of research on the role of water flux as an instructive factor in patterning cell identity in meristems and beyond. While the example of the deep boundary may appear as an extreme case due to strong tissue curvature, we believe that such shrinkage might be more widespread. For instance, we could observe this effect in the region below the growing organs in non-dissected shoot apices. Such water fluxes may in turn have many impacts on patterning tissue growth and identity. For instance, it has recently been shown that the inhibition of lateral roots during xerobranching response to dry soil, is controlled by a change in internal fluxes that reshapes hormonal gradients (32). It is thus possible that, despite being less detectable, water fluxes contribute to patterning of many more tissues in plants, and likely in other kingdoms too. For instance, osmolarity can impact membrane tension and cell polarity, with shared mechanisms between plants and animals (33). One could imagine that water fluxes also contribute to tissue patterning in animal embryos.

## Acknowledgments

We thank the Mosaic and MechanoDevo teams at the RDP lab for insightful discussion, and Platim for help with imaging. We thank Johanna Dickmann and Marianne Lang for helpful comments on this manuscript.

## Funding

This work was supported by an EMBO Long Term Fellowship (#ALTF1-2020 to J.A-S.) and by the European Research Council (ERC-2021-AdG-101019515 “Musix” to O.H.)

### Author contributions

Conceptualization: JAS, IC, CG, OH

Methodology: JAS, IC, AK, GC, CL

Funding acquisition: JAS, OH

Project administration: CG, OH

Supervision: CG, OH

Writing – original draft: JAS, IC, OH

Writing – review & editing: JAS, IC, AK, GC, CG, OH

## Data and materials availability

All data are available on demand and/or in the main text and supplementary material.

## Supplementary Materials

Materials and Methods

Supplementary references

Figs. S1 to S5

Data S1

