## Supplementary data for "Water fluxes contribute to growth patterning in shoot meristems"

### Growth-derived water fluxes define the meristem boundary

**The PDF file includes:**

Material and Methods
Supplementary references
Figs. S1 to S5

**Other Supplementary Materials for this manuscript include the following:**

Data S1 (Excel file)

### Materials and Methods

Plant materials and growth conditions

Plasma membrane marker *pUBQ10::LTi6B-TdTomato* (*2*) has been previously described. Dissected meristems and naked meristems (without organs) were obtained and processed as previously described (*6*). Briefly, two protocols were used to image shoot apical meristems:

(i) For analysis of reproductive meristems plants were grown on soil for 4 weeks under long-day (16 hours light / 8 hours dark) conditions. Reproductive meristems were dissected right after bolting when the stem was 1-2 cm. After dissection reproductive meristems were transferred to the previously described apex culture medium (ACM) supplemented with N6-benzyladenine (500nM) (*6*). Shoot apices were kept in a long day chamber until and during imaging.

(ii) Naked meristems were obtained by growing sterilized seeds in Arabidopsis media containing 10µM of NPA (N-1-Naphthylphthalamic Acid) (Sigma) as previously described (*6*), pin-like meristems emerged after 3 to 4 weeks depending on the line. The plants were then transferred to NPA-free medium to induce organ outgrowth (most experiments), or was left on NPA medium to maintain a naked meristem.

HPTS incubation experiments

Inflorescence meristems from long day grown plants were used in all HPTS incubations. After bolting, 5 cm - tall stems were cut at the base and immediately transferred to an Eppendorf tube containing liquid ACM (without agarose) supplemented with 2.5 mg/mL of HPTS (Molecular Probes, Leiden, Netherlands). Tubes for control samples contained only liquid ACM. Stems were then incubated in a humid chamber (i.e. saturated hygrometry with a wet paper) under long day conditions for 1 (n=5), 3 (n=5), 6 (n=10), 12 (n=10), and 24 hours (n=5). Stems were dissected right after the incubation time and placed in a Petri dish with solid ACM for imaging within the following 30 mins. In order to obtained the profile of HPTS signal intensity in L1 and L2, we used ImageJ (Z project plugin) to generate maximum intensity projections of 10 µm thick orthogonal image stack of different primordium stages (P5 and P7). Next a region of interest was selected using a line (10 µm thick) that covers the L1 and L2 cell layers from the central zone to the abaxial side of primordium. Signal intensity levels for 5 meristems from 5 independent plants where then plotted using the online tool Plot of Twist (https://huygens.science.uva.nl/PlotTwist/). HPTS experiments with NPA-induced pin-like meristems were not possible because root or leaf-provided HPTS failed to reach the shoot apex.

Model description

We built an abstract bidimensional multicellular description of a SAM in the framework of the so-called vertex-based models (*10*) where pressure in the cells equilibrates with tension in the walls. As in the Lockhart model (*11*), this mechanical equilibrium is coupled to the growth of the walls and to osmosis-based water fluxes between cells and between cells and water sources.

*Mechanical equilibrium:* Let $P_i$ be the turgor pressure in each cell $i$, $\varepsilon_k^e$ the elastic deformation of each wall $k$, and $E_k$ its elastic modulus ($\sigma_k = E_k\varepsilon_k^e$ is then the mechanical stress). The tissue being at every moment in quasi-static equilibrium, pressure forces on wall edges and elastic forces within walls balance exactly; in a vertex-based model, this leads to, at each vertex $v$:

$$\frac{1}{2}\sum_{k\sim v}\Delta_k P A_k \boldsymbol{n}_k + \sum_{k\sim v} E_k\varepsilon_k^e a_k \boldsymbol{e}_{k,v} = 0$$

Where the sums are computed over the indices of walls $k\sim v$ adjacent to vertex $v$, and $\Delta_k P = P_{k_1} - P_{k_2}$ is the pressure jump across wall face $k$, $k_1 < k_2$ are the indices of the cells separated by face $k$, $A_k$ is the area of the face $k$ on which pressure is exerted, $\boldsymbol{n}_k$ is the normal vector to face $k$, oriented from cell $k_1$ to cell $k_2$, $a_k$ is the cross-section of the face, on which the elastic stress is exerted, and, finally, $\boldsymbol{e}_{k,v}$ is the unit vector in the direction of face $k$, oriented from vertex $v$ to the other end of face $k$. In the case of the geometry of the Lockhart model, this mechanical equilibrium leads to a constant proportionality between pressure and wall deformation, but in the more general case here, the relation between pressure and wall deformation depends on the varying geometry of the cells.

*Cell wall growth:* The cell walls are modeled as a visco-elasto-plastic material, similarly to the Ortega model (*12*). Let $l_k$ be the length of wall $k$, the rate of change of $\varepsilon_k^e$ is given by:

$$\frac{d\varepsilon_k^e}{dt} + \phi_k^w E_k(\varepsilon_k^e - \varepsilon_k^Y)_+ = \frac{1}{l_k}\frac{dl_k}{dt}$$

Where $\phi_k^w$ is the extensibility and $\varepsilon_k^Y$ is the yield deformation of wall $k$, and $(x)_+$ is the positive part of any real number $x$. This equation describes the fact that a wall $k$ with elongation rate $\frac{1}{l_k}dl_k/dt$ (*e.g* because of an influx of water) will develop an elastic deformation and therefore tension, and that growth will relax any elastic deformation above the threshold $\varepsilon_k^Y$ with a characteristic time $1/(\phi_k^w E_k)$.

*Water fluxes:* In the context of plant tissues (*13*), the water potential reduces to $\Psi = P - \pi$, where $\pi$ is the osmotic pressure. As in the Lockhart model, fluxes across a perfectly semi-permeable membrane of surface $A$ and permeability $L$ follow decreasing water potential gradients and are in the form $U = AL\Delta\Psi$. In the model presented here, cells can exchange water between them and with the apoplasm which is assumed to have a constant water potential $\Psi^a = 0$, and behaves as a water supply for the growth of the cells. In addition, we assume that water fluxes account for all the volume variations of the cells, and we get, for each cell $i$:

$$\frac{dV_i}{dt} = A_i L_i^a (\Psi_i - \Psi^a) + \sum_{j\sim i} A_{ij} L_{ij}^s (\Psi_j - \Psi_i)$$

Where $A_i$ is the surface of cell $i$, $L_i^a$ its permeability with the apoplasm, $\Psi_i$ its water potential; the sum is performed over the neighbor cells $j\sim i$ adjacent to cell $i$, and $A_{ij}$ is the common surface between cells $i$ and $j$, and $L_{ij}^s$ the water permeability. For parsimony, the osmotic pressure is assumed constant and homogeneous in the cells.

The coupling between these three equations results in a complex mathematical and computational problem for which we provided a numerical resolution in (*14*). Pressure and growth rate are not prescribed in this model, as they emerge from the interaction between fluxes and wall rheology, through the mechanical equilibrium. This model has already been quantitatively compared to biological data and predicted pressure and growth rate heterogeneities in the SAM (*15*).

*Simulations presented in this article:* The goal here is to represent the differential growth between organs and the central zone in the SAM (Fig. S3), with water supplied from below. In order to avoid boundary conditions between the SAM and the tissue underneath, we choose an initial circular geometry with 316 cells organized in concentric layers (Fig. S3A). This bidimensional mesh can be interpreted as a section through the SAM along the axis of the stem, as drawn in Fig. 3J. Then, the identity of the cells is determined by the choice of the parameters of the model (Fig S3A and Table S1). Cells in the center, below L3, do not grow ($\phi^w = 0$) and are connected to a water source ($L^a \neq 0$), they mimic the basal part of the SAM that is connected to vascular bundles providing water. Conversely, cells in L1-L3 layers are isolated from the water source ($L^a$ is three orders of magnitude lower) and grow thanks to the water from cells below; water mostly travels from cell to cell in L1-L3 which implies that they are in competition for water. The elastic modulus in L1-L3 (resp. outer walls) is 75% of cells below (resp. 150%). Starting from an initial state with zero elastic deformation, the tissue is put under pressure until the center cells have reached a stationary pressure, and cells in L1-L3 have started their plastic growth. This state is reached after 6.64 h in the simulation, and this marks the time $t = 0$. The detailed values of the parameters are given in Fig. S3A and Table S1. The outgrowth of the primordium is then triggered by a further decrease of the elastic modulus in a few cells in L1-L3 (x 0.75 in the reference simulation, x 0.5 and x 0.75 in the parameter exploration below, Fig. S4).

This abrupt decrease of the elastic modulus induces 1) a global drop of the pressure (Fig. S3C), especially in the primordium, and 2) a peak in the growth rate of the primordium and a drop in the boundary cells B1, B2, B3 (Fig. S3E). This initial fast response corresponds to a transient elastic response, and after $\approx$ 1h the cells have adapted to the change of conditions. As explained in the main text, the boundary cells B3 end up losing water to the primordium. Note that this shrinkage is not due to compression as the pressure in the primordia is lower than the pressure in the boundary. The pressure and growth rate are globally decreasing with time, as an effect of the global increase of cell volume, as analyzed in (*14*). One can also notice an apparently periodic growth rate heterogeneity in the region far from the primordium, this is due to a toplogy (number of neighbours) heterogeneity in the construction of the mesh, as analyzed in (*15*).

**Table S1.** Values of the parameters before the organ initiation

| **Parameter** | **Description** | **Quantity** |
|---|---|---|
| $\varepsilon^Y$ | Threshold elastic deformation | 5% |
| $E_0$ below L3 | Cell wall elastic modulus | 90 MPa |
| $E$ in L1, L2 and L3 (initial) | Cell wall elastic modulus | $0.75E_0$ |
| $E$ in outer periclinal wall | Cell wall elastic modulus | $1.5E_0$ |
| $\phi^w$ in L1-L3 | Cell wall extensibility | $5.5 \times 10^{-5}$ $MPa^{-1}.s^{-1}$ |
| $\phi^w$ below L3 | Cell wall extensibility | 0 $MPa^{-1}.s^{-1}$ |
| $L_0^s$ below L3 | Water conductivity between cells | $3 \times 10^{-9}$ $m.MPa^{-1}.s^{-1}$ |
| $L^s$ in L1 and L2 | Water conductivity between cells | 5 x $L_0^s$ |
| $L^s$ in L3 | Water conductivity between cells | 10 x $L_0^s$ |
| $L_0^a$ below L3 | Water conductivity with external source | $1.56 \times 10^{-10}$ $m.MPa^{-1}.s^{-1}$ |
| $L^a$ in L1, L2 and L3 | Water conductivity with external source | $L_0^a/1000$ |
| $\pi$ | Osmotic pressure | 1 MPa |

<u>Image analysis</u>

*Image acquisitions*

All confocal acquisitions were performed using a Leica SP8 or a Zeiss LSM980 microscope. A SP8 microscope with a resonant scanner was used for the time-lapse acquisitions on dissected meristems and pin-like meristems were imaged with Zeiss LSM980.

*Volumetric analysis*

In order to improve the quality of the plasma membrane marker signal, all confocal stacks were resampled to get isometric voxels and then processed by an anisotropic diffusion filter (*15*). Then, to increase the signal level in the outer periclinal membranes of the epidermal cell layer, we used a Level Set Method to locate the tissue surface precisely (*16*) and added the obtained contour to the original image (Fig. S6A). This step was particularly important to accurately delimit cells in the boundary region of the meristem.

Then we performed an automated 3D seeded watershed segmentation derived from the MARS algorithm (*17*) and obtained a 3D labelled image stack (Fig. S6B). Using the Blockmatching method (*18*), we registered the membrane marker images of consecutive time points to obtain the 3D vector field deforming the tissue at T0 onto the one a T0+T. We applied this geometrical transformation to the segmented image at T0, and automatically extracted the cell lineages by identifying the most overlapping cells between T0 and T0+T. These cell lineages allowed us to directly evidence new cell divisions (Fig. S6C). We used the implementations of the segmentation and registration algorithms provided in the Tissue Image ToolKit Python library (https://mosaic.gitlabpages.inria.fr/timagetk).

We quantified morphological cell properties such as cell volumes directly on the 3D labelled image. Then to estimate the mean curvature at the level of epidermal cells (Fig. S6D), we extracted a triangle mesh of the tissue surface, using decimation and smoothing on the mesh obtained by applying the Marching Cubes method on the binary mask of the segmented image. The surface mesh extraction was performed using the implementations provided in the VTK library (*19*). We computed the principal curvatures at the level of each triangle of this mesh based on the vertex normal vectors (*20*) and averaged this information on the mesh vertices (area-weighted average of incident triangles). The mean curvature of the tissue surface at the level of a given epidermal cell was then estimated as the mean curvature at the vertex of the surface mesh that lies closest to its center. Cell sphericity was calculated using the following formula: Sphericity = $36\,\pi * V^2/S^3$, where V is the cell volume and S is the total cell surface.

Automated scripts performing these successive image analysis steps, as well as visualization of the output segmentations and quantified properties, were made available in the titk_tools Python package (https://gitlab.inria.fr/gcerutti/boundary_registration).

Since the XY resolution (ca. 200 nm) in our images was different from that in the Z orientation (ca. 500 nm), we aimed at identifying the error in volumetric measurements by comparing multiangle acquisitions in pin-like meristems. To do so we imaged meristems first in the standard vertical position (VP1) then in a tilted angle of 30° (TA), and finally in the vertical position again

(VP2). The time between confocal acquisitions was less than 2 mins. The error associated to consecutive confocal imaging (VP2 -VP1) was 5.5 ± 3%. The error associated to the Z distortion (VP1-TA) – (VP1-VP2) = 0.27± 5%.

All box plots were generated using the online tool Plots of Data (https://huygens.science.uva.nl/PlotsOfData/)

http://ieeexplore.ieee.org/document/1348359/), pp. 288–297.

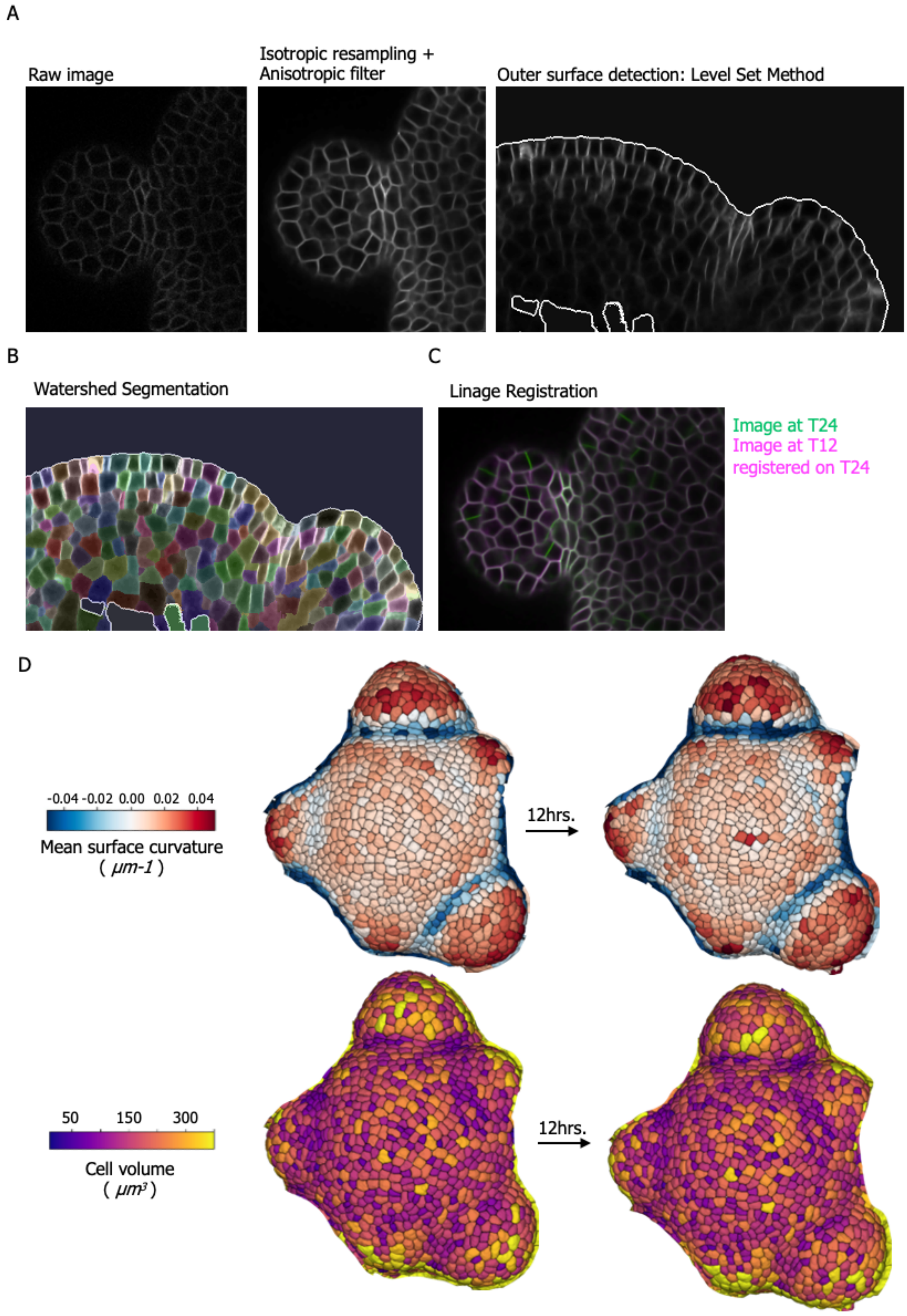

**Fig. S1.** Processing pipeline for 3D segmentation

(A) Data preprocessing consisted of applying isotropic resampling and anisotropic filtering to improve the signal of the plasma membrane marker. In addition, signal from the outer anticlinal membranes was enhanced using a Level Set Method to accurately segment central zone and boundaries. (B) Image after watershed segmentation with individual cell IDs indicated by colors. Cell lineages were obtained by registering the stack obtained in T12 (magenta) on the stack obtained in T24 (green). (C) New cell divisions are visible in the green channel. (D) Heatmaps representing cell properties in the same meristem before and after 12 hours of growth.

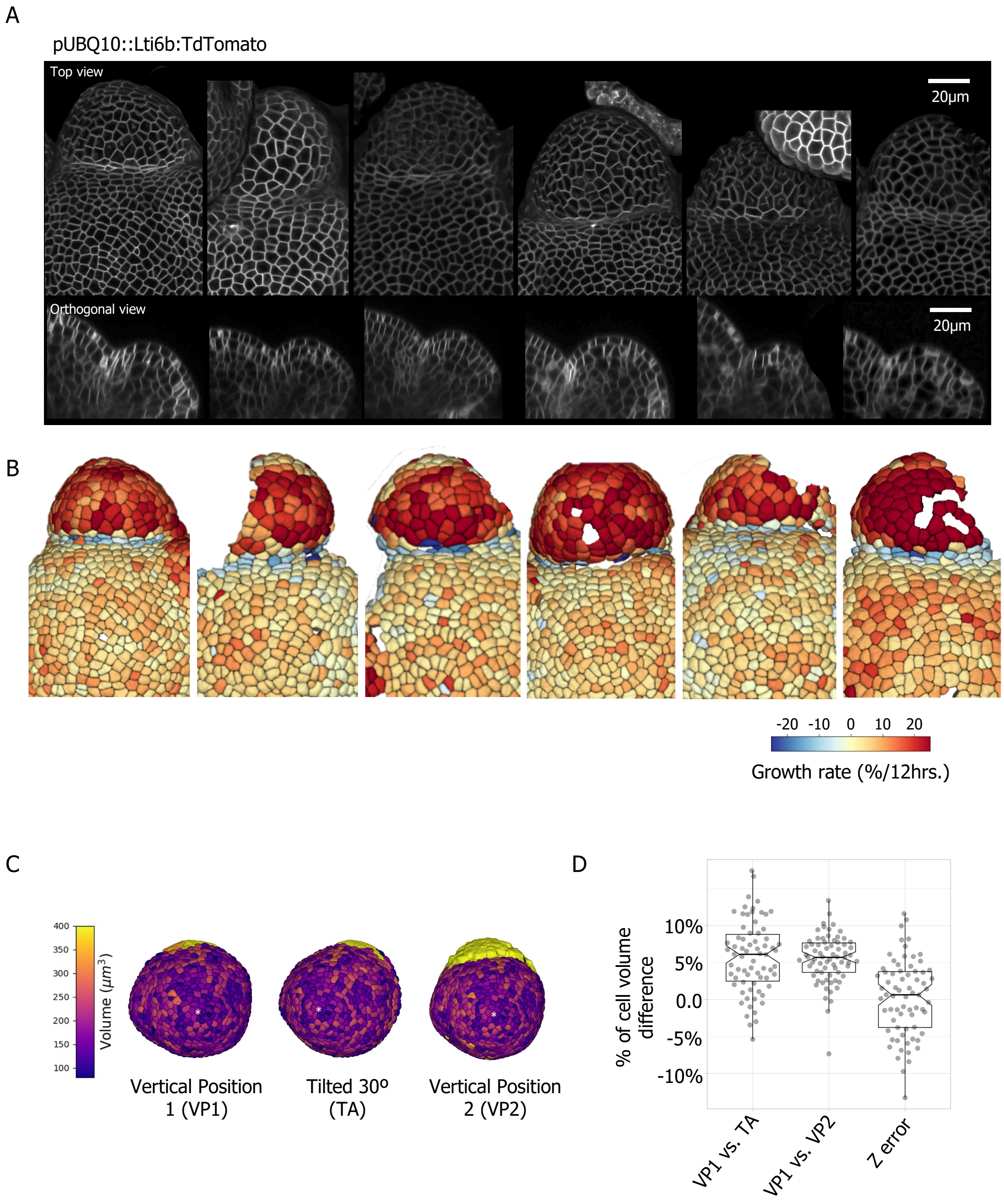


**Fig. S2.** Quantitative analysis of cell volume change in dissected and pin-like meristems.

(A) Processed plasma membrane signal (L1 only signal) from 6 dissected meristems originating from 6 independent plants. Below, an orthogonal view was obtained from the P5 primordium which in all cases revealed boundary shrinkage. (B) Volumetric growth rate from the 6 samples shown in (A). (C) Multi-angle comparison of cell volumes in pin-like meristem. The first acquisition was obtained in vertical position 1 (VP1). Afterwards, the sample was tilted (Tilted acquisition (TA)) and brought back to the original position (vertical position 2 (VP2)). The white asterisk indicates the same cell across imaging. (D) Measurements of the volume of 70 cells compared across acquisitions shows that multiple immediate imaging (in less than 2 min) leads to a reduction in cell volume by ~ 6%. Therefore, to identify the Z distortion (Z error) we differentiate the effect of consecutive imaging from the effect of sample orientation (see methods for details).

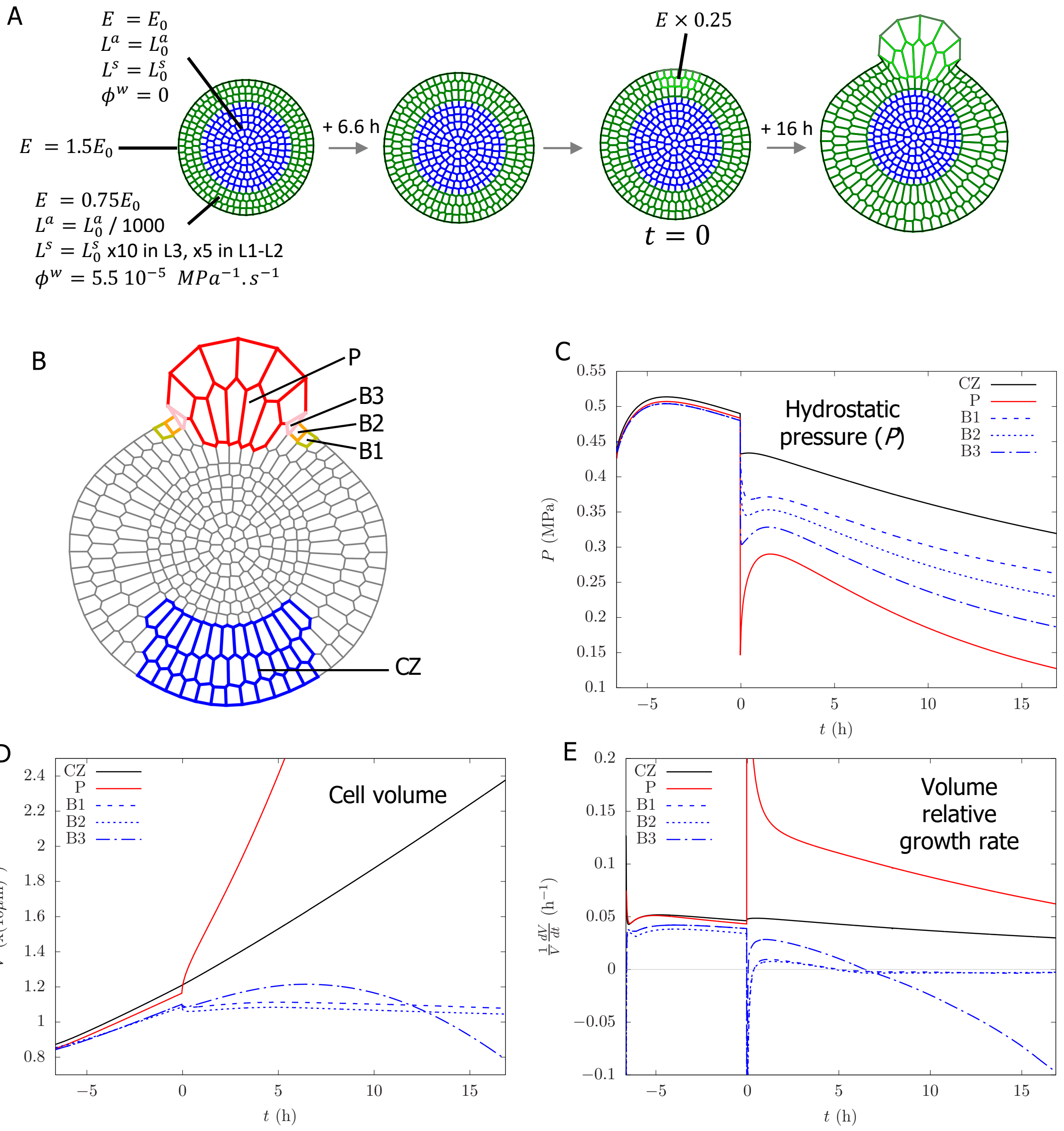


**Fig. S3.** Time evolution of the reference simulation.

(A) Scenario presented in the main text: initiation with a zero elastic deformation with regions defined by heterogeneous parameters; initial pressurization of the tissue; initiation of the primordium at $\boldsymbol{t = 0}$; growth of the primordium. (B) Regions where the time evolution is monitored: mean values over primordium cells (P) and central zone (CZ), individual cell values for the boundary cells (B1, B2 and B3). (C-E) Time evolution of hydrostatic pressure (B), cell volume (D) and volume relative growth rate (E). Time $\boldsymbol{t = 0}$ indicates the initiation of the primordium with a reduction of the elastic modulus.

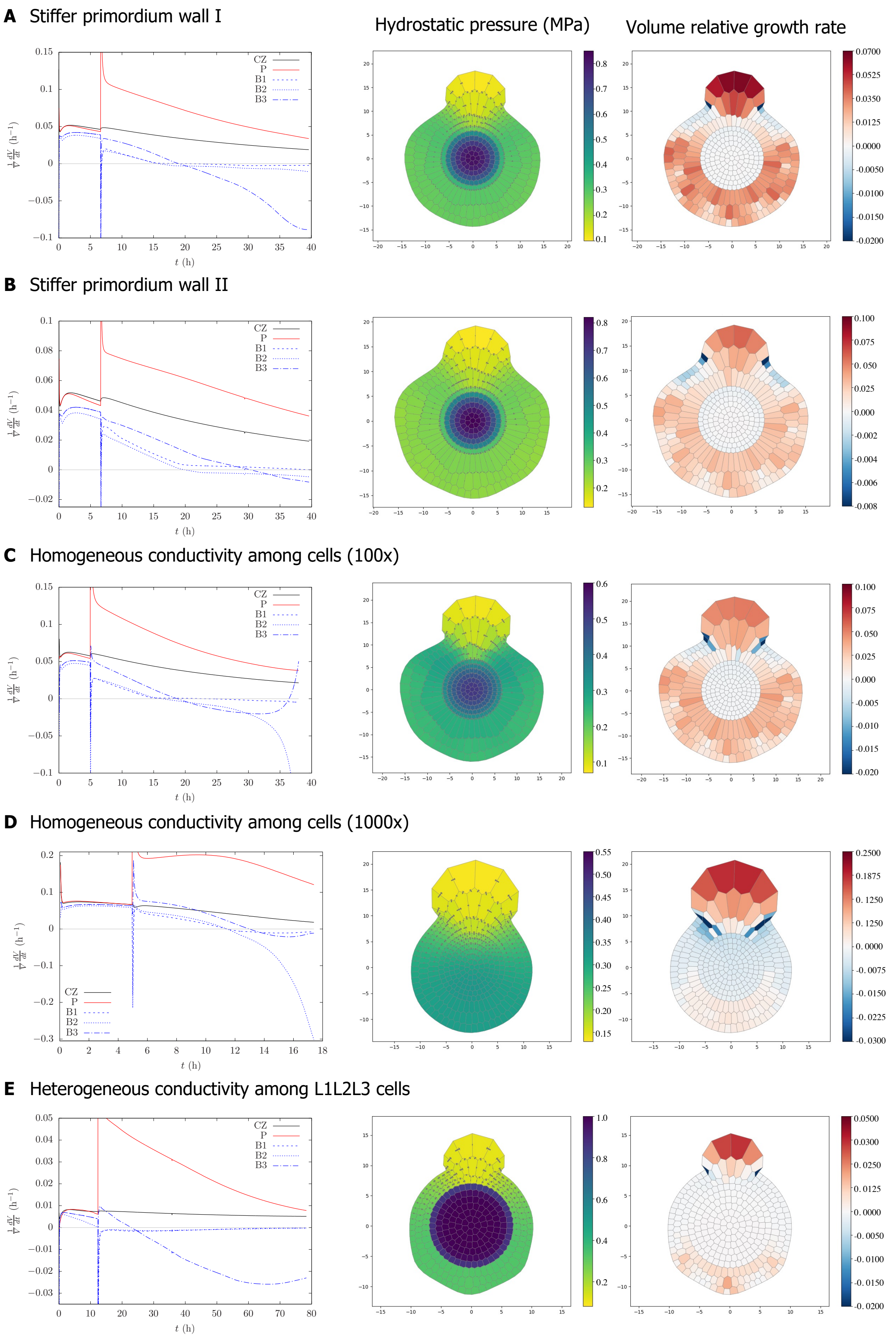
A Stiffer primordium wall I
Hydrostatic pressure (MPa)
Volume relative growth rate
B Stiffer primordium wall II
C Homogeneous conductivity among cells (100x)
D Homogeneous conductivity among cells (1000x)
E Heterogeneous conductivity among L1L2L3 cells
CZ
P
B1
B2
B3
t (h)
$\frac{1}{V}\frac{dV}{dt}$ (h$^{-1}$)

**Fig. S4.** Effect of parameter variation in the model

(A-E) Plots of volume relative growth rates and heatmaps representing the final hydrostatic pressure (green-blue) and final volume relative growth rates (red-blue) after simulations with different parameter conditions. A stiffer wall in emerging primordia affects growth rates and the topology of the mesh. Higher cell-to-cell conductivity additionally results in cell shrinkage in inner layers L2 or L2L3. No difference in L1-L3 cell-to-cell conductivity results in lower positive growth rates and cell shrinkage in only one pair of boundary cells. A) Stiffer primordium wall I: Elastic modulus $E$ (primordium creation) = 34 Mpa. B) Stiffer primordium wall II: Elastic modulus $E$ (primordium creation) = 50,6 Mpa. C) Homogeneous conductivity among cells (100x more): $L^s$= 3 $10^{-7}$ m.MPa$^{-1}$.s$^{-1}$ and Elastic modulus $E$ (primordium creation) = 34 Mpa. D) Homogeneous conductivity among cells (1000x more): $L^s$= 3 $10^{-6}$ m.Mpa$^{-1}$.s$^{-1}$ and Elastic modulus $E$ (primordium creation) = 34 Mpa. E) Heterogeneous conductivity among L1L2L3 cells: $L^s$ (L1L2L3)= 3 $10^{-7}$ m.MPa$^{-1}$.s$^{-1}$, $L^s$ (other cells)= 3 $10^{-9}$ m.MPa$^{-1}$.s$^{-1}$ and Elastic modulus $E$ (primordium creation) = 34 Mpa.

A

| GO Term | Observed Frequency | *p*-value |
|---|---|---|
| translation | 52 out of 506 genes, 10.3% | 3,15E-19 |
| macromolecule biosynthetic process | 64 out of 506 genes, 12.6% | 3,96E-06 |
| cellular macromolecule biosynthetic process | 63 out of 506 genes, 12.5% | 4,67E-06 |
| pattern specification process | 15 out of 506 genes, 3% | 1,08E-05 |
| developmental process | 69 out of 506 genes, 13.6% | 8,43E-05 |
| reproductive process | 42 out of 506 genes, 8.3% | 0,000161 |
| **response to salt stress** | 25 out of 506 genes, 4.9% | 0,000177 |
| **response to osmotic stress** | 25 out of 506 genes, 4.9% | 0,000485 |
| cellular component biogenesis | 15 out of 506 genes, 3% | 0,000551 |
| cellular component biogenesis at cellular level | 15 out of 506 genes, 3% | 0,000551 |

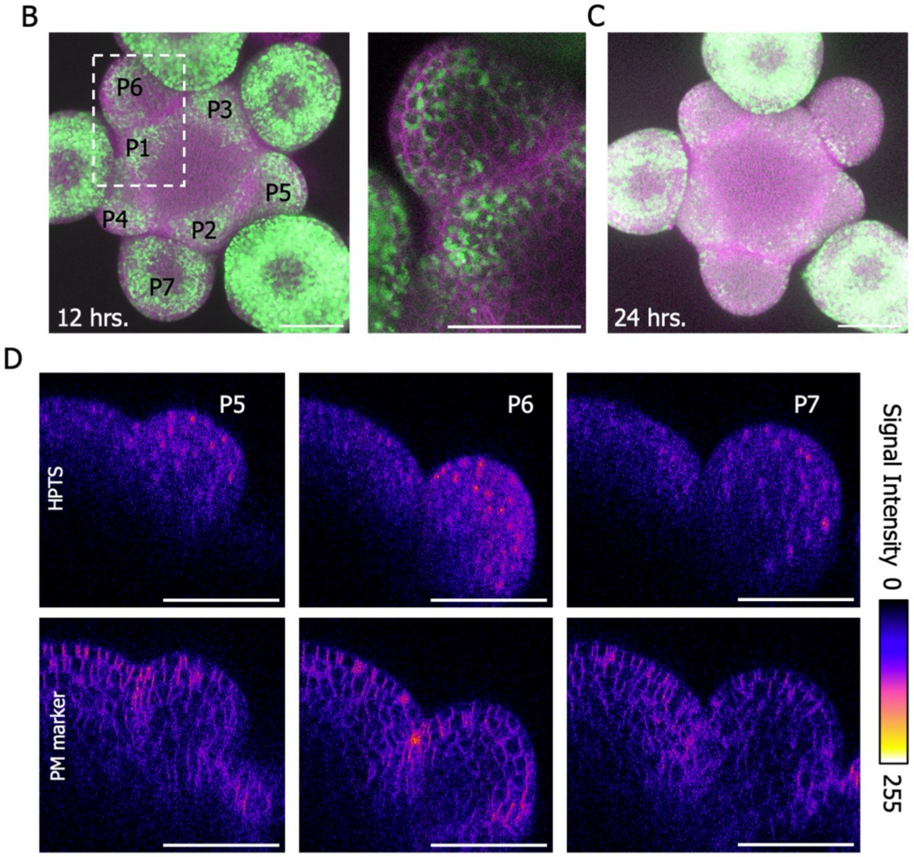


**Fig. S5.** HPTS allocations pattern in the SAM

(A) Top ten enriched GO terms in a list of 506 boundary-specific upregulated genes. (B) Maximum intensity projection of HPTS pattern with the plasma membrane marker after 12. of incubation. A transversal slice perpendicular to the stem and in the middle of the meristem was obtained from the white inset box in the 12 hours incubation image to show the apparent vacuolar localization of HPTS. (C) Maximum intensity projection of HPTS pattern with the plasma membrane marker after 24 hours of incubation. (D) Orthogonal sections showing HPTS and plasma membrane (*pUBQ10::LTi6B-TdTomato*) signal intensities for organs P5 to P7 after 6 hours of incubation (stacks from Fig. 4C).

**Data S1 (Excel file).** GO enrichment analysis on upregulated genes from SAM domains.
